# Epigenetic rejuvenation of the hippocampus by environmental enrichment

**DOI:** 10.1101/776310

**Authors:** Sara Zocher, Rupert W. Overall, Mathias Lesche, Andreas Dahl, Gerd Kempermann

## Abstract

The decline of brain function during aging is associated with epigenetic changes, including DNA methylation. Lifestyle interventions can improve brain function during aging, but their influence on age-related epigenetic changes is unknown. Using genome-wide DNA methylation sequencing, we here show that environmental enrichment counteracted age-related DNA methylation changes in the hippocampal dentate gyrus of mice. Specifically, environmental enrichment prevented the aging-induced CpG hypomethylation at target sites of the methyl-CpG-binding protein Mecp2, which is known to control neuronal functions. The genes at which environmental enrichment counteracted aging effects have described roles in neuronal plasticity, neuronal cell communication and adult hippocampal neurogenesis and are dysregulated with age-related cognitive decline in the human brain. Our results highlight the rejuvenating effects of environmental enrichment at the level of DNA methylation and give molecular insights into the specific aspects of brain aging that can be counteracted by lifestyle interventions.

## Introduction

Aging is associated with a progressive decline in brain function that manifests in cognitive impairments, increased risk for neurodegenerative diseases and loss of neural plasticity. Lifestyle factors, including physical exercise and cognitive stimulation, attenuate age-related reductions of brain function in humans^1^, contributing to what is called ‘reserve’ or ‘maintenance’ of brain function^2, 3^. In rodent models, environmental enrichment (ENR) promotes life-long brain plasticity and health^4, 5^. ENR stimulates the function of existing neurons and adult neurogenesis in the hippocampus—a brain region with key roles in learning and memory, but high susceptibility to stress- and age-related impairments^6^. We set out to investigate the molecular mechanisms that link environmentally induced brain plasticity with improved brain maintenance during aging.

Genome-wide DNA methylation changes are hallmarks of cellular aging^5^ and are used as biomarkers of the aging process^6^. In the hippocampus, the aging-induced dysregulation of the DNA methylation machinery is involved in the development of cognitive impairments^9, 10^. Proper control of neuronal DNA methylation patterns in the adult brain is crucial for dynamic gene expression changes associated with synaptic plasticity and memory formation^11–13^. The sensitivity of neuronal methylomes to environmental stimuli, such as maternal behaviour^14^, learning stimuli^15^ or isolated neuronal activation^16^, contributes to the molecular mechanisms underlying experience-dependent brain plasticity^17^. Conversely, aberrant changes of neuronal DNA methylation patterns have been described in aging^18, 19^ and in age-related disorders, such as Alzheimer’s disease^20, 21^. Since neuronal DNA methylation patterns are plastic and sensitive to environmental experiences, behavioral interventions could potentially rescue such aberrant DNA methylation changes and thereby promote brain health in old age.

We therefore investigated the influence of life-long ENR on DNA methylation patterns in the hippocampal dentate gyrus and found that ENR counteracted aging-induced DNA methylation changes at genes related to neuronal plasticity and adult hippocampal neurogenesis. Our results highlight the potential of lifetime experiences to influence brain health in old age and provide a possible mechanism underlying the effects of lifestyle factors on brain aging.

## Results

### Environmental enrichment changes DNA methylation at brain plasticity genes in the adult dentate gyrus

To first investigate whether ENR changes DNA methylation patterns in the adult dentate gyrus, female C57BL/6JRj mice were kept in ENR or standard housing cages (STD) for three months starting at an age of six weeks. Genome-wide DNA methylation profiling was performed on micro-dissected dentate gyrus tissue by reduced representation bisulfite sequencing (RRBS)^22, 23^. No ENR-induced changes in global CpG methylation levels were detected (Fig. 1a), underscoring general genomic stability after ENR. Nevertheless, ENR modified methylation levels at 11,101 individual CpGs, i.e. 1.25 % of all covered CpGs (Fig. 1b). Differentially methylated CpGs (dmCpGs) were depleted at CpG islands, CpG island shores, promoters and exons but enriched at enhancers, introns and intergenic regions of the genome (Fig. 1c). Additionally, ENR changed methylation at 0.019 % of CpHs (in total only 750 differentially methylated CpHs; dmCpHs), which showed a similar genomic distribution to dmCpGs but no enrichment at enhancers (Supplementary Fig. 1; Supplementary Data 1). Thus, ENR changed methylation in the dentate gyrus predominantly at CpGs located within regulatory genomic regions.

**Fig. 1:**
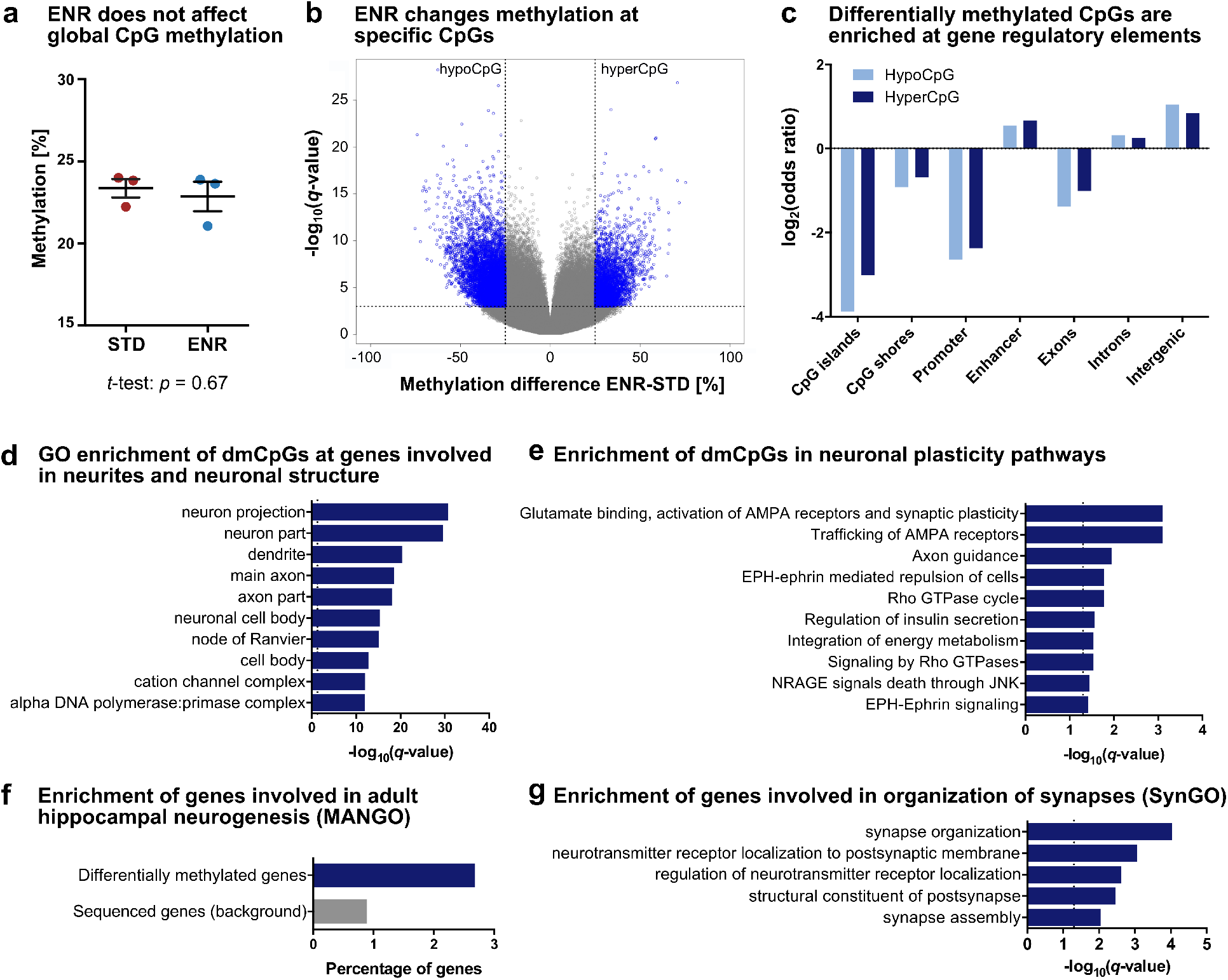
Environmental enrichment changes CpG methylation in the adult dentate gyrus at genes with known role in neuronal plasticity, hippocampal neurogenesis and synapse organization. DNA methylation differences between 4.5-month-old mice housed in environmental enrichment (ENR) for three months and age-matched standard housed mice (STD) (n = 3 per group). **a**, ENR does not change global CpG methylation. **b**, Volcano plot highlighting ENR-induced differentially methylated CpGs (dmCpGs; methylation difference > 25 % and *q* < 0.001) in blue. Among the dmCpGs, 67.6 % decreased methylation (hypoCpG) and 32.4 % increased methylation in ENR (hyperCpG). **c**, Genomic distribution of dmCpGs. HypoCpG and hyperCpG are depleted at CpG islands, CpG island shores, promoters and exons (log_2_(odds ratio) < 0) and enriched at enhancers, introns and intergenic regions (log_2_(odds ratio) > 0). Adjusted *p*-values < 0.001 for all regions. **d**, Top ten significantly enriched cellular components from Gene Ontology (GO) enrichment analysis with differentially methylated genes (373 genes). **e**, ENR changed CpG methylation at genes involved in synaptic plasticity. Depicted are the top ten significantly enriched pathways from the Reactome database. **f**, Genes functionally involved in hippocampal neurogenesis (as annotated in the MANGO database) are enriched among ENR-induced differentially methylated genes. Hypergeometric test: *p* = 0.0020. **g**, Differentially methylated genes are involved in synapse organization and signaling. Depicted are significantly enriched biological processes from the SynGO database. The term “regulation of neurotransmitter receptor localization” was shortened from “regulation of neurotransmitter receptor localization to postsynaptic specialization membrane”. Dashed lines in (d-e, g) indicate significance thresholds of *q* < 0.05.

To explore the neuronal processes that are regulated by ENR, we performed gene set enrichment analysis with the 373 ENR-induced differentially methylated genes using expert-curated knowledge bases. Gene ontology (GO)^24^ and Reactome pathway analyses^25^ showed that ENR-induced dmCpGs were enriched at genes involved in structural components of neurons, such as “axon part”, “dendrite” and “dendritic spine” (Fig. 1d), and at genes with known functions in synaptic plasticity pathways, including glutamate receptor signaling and axon guidance (Fig. 1e; Supplementary Data 2). Enrichment analysis for genes from the Mammalian Adult Neurogenesis Gene Ontology (MANGO)^26^ highlighted that ENR changed DNA methylation at genes with described function in hippocampal neurogenesis, such as *Fgfr1*, *Gria1*, *Nfatc4*, *Ntf3*, *Flt1* and *Thrb* (Fig. 1f). In addition, enrichment analysis using the Synaptic Gene Ontologies (SynGO) knowledgebase^27^ suggested that ENR-induced differentially methylated genes are enriched at genes involved in synaptic assembly, organization of post-synapses and neurotransmitter signaling (Fig. 1g). These results indicated that ENR regulates pathways involved in neuronal plasticity and adult hippocampal neurogenesis in the murine dentate gyrus.

### Age-related DNA methylation changes in the dentate gyrus

To investigate the influence of ENR on age-related DNA methylation changes in the dentate gyrus, RRBS was performed on dentate gyrus tissue from young (6.5-week-old) and aged (14-month-old) mice, which had lived in STD or ENR for four days (young) or over a year (aged).

To determine age-related DNA methylation changes in the dentate gyrus, we compared DNA methylation profiles between young STD mice and aged STD mice and detected 41,961 dmCpGs and 6,600 dmCpHs (Supplementary Fig. 2a–b; Supplementary Data 3). While most dmCpGs were hypomethylated in aged mice (77.93 % of dmCpGs), aging predominantly increased methylation of CpHs (74.05 % of dmCpHs). The gene locations of aging-induced methylation changes in the dentate gyrus were significantly enriched with genes previously reported to exhibit age-related methylation changes in different tissues (Supplementary Fig. 2c). Additionally, more than 92.0 % of the here identified age-related differentially methylated genes have been shown to change methylation during aging in the mouse hippocampus by two independent studies^18, 28^, highlighting the robustness of aging-associated DNA methylation changes in the brain.

We found that age-related differentially methylated genes were significantly enriched in pathways related to neuronal plasticity, neuronal signaling and energy metabolism (Supplementary Fig. 2d; Supplementary Data 4). Remarkably, the highest enriched pathways of genes with age-related DNA methylation changes overlapped considerably with the highest enriched pathways of ENR-induced differentially methylated genes in the non-aged brain (compare Fig. 1e), which suggested that aging and ENR changed DNA methylation at genes involved in similar pathways.

### ENR counteracts age-related DNA methylation changes in the dentate gyrus

To investigate the effect of life-long ENR on aging, we first compared global aging-induced DNA methylation changes between STD and ENR mice. In STD mice, aging was associated with a global 13.81 % decrease in CpG methylation (Fig. 2a), which is in accordance with the predominant CpG hypomethylation found at the individual CpG level (compare Supplementary Fig. 2). In contrast, aged ENR mice did not show significant global CpG methylation differences compared to young STD mice, suggesting that the age-related global CpG hypomethylation in the dentate gyrus is at least partially prevented by ENR. No ENR- and age-related global methylation changes were seen in the CpH context (Supplementary Fig. 3).

**Fig. 2:**
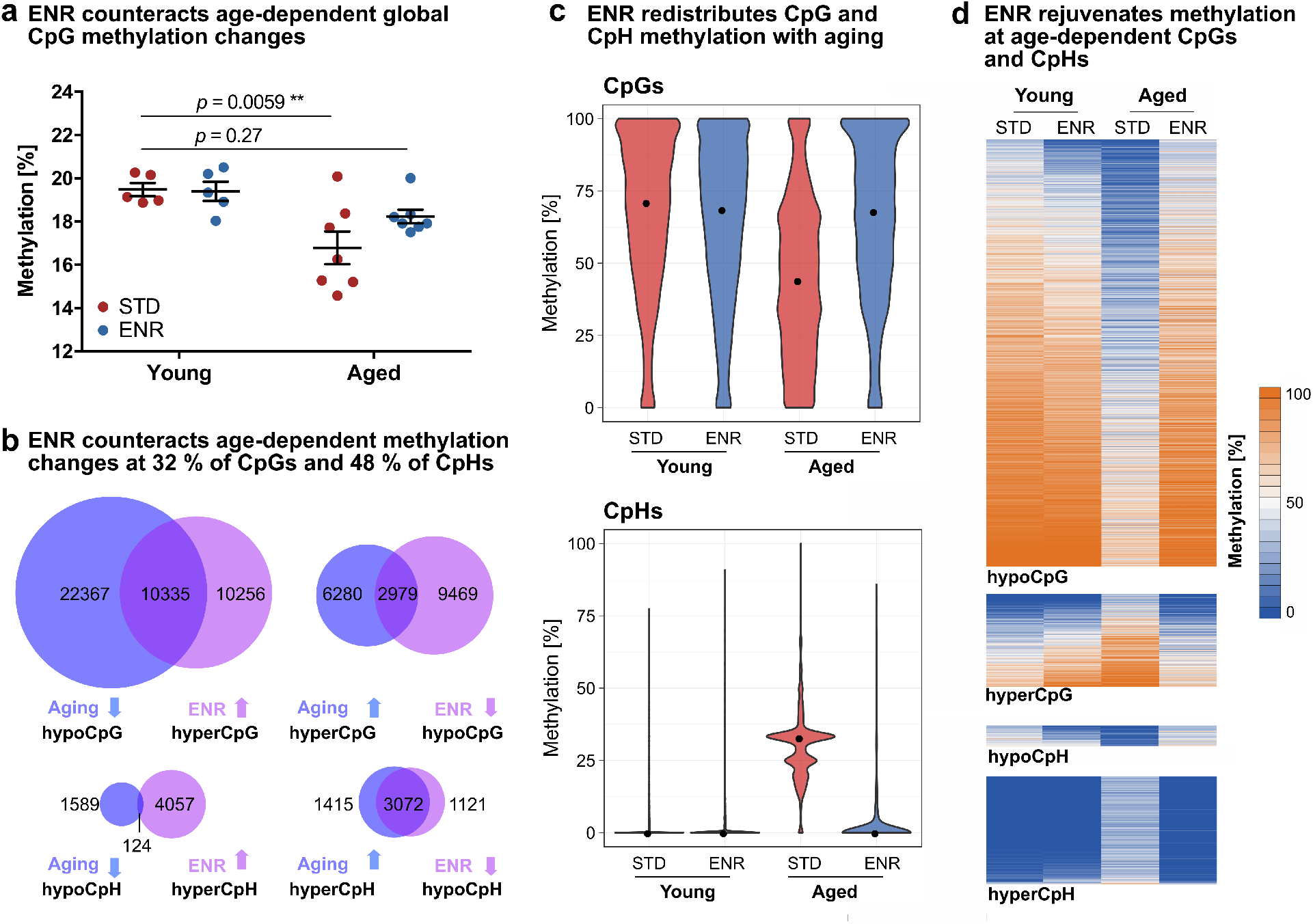
Environmental enrichment protects the dentate gyrus from age-related CpG and CpH methylation changes. DNA methylation profiles of young mice housed in ENR or STD for four days (Young ENR; Young STD; n = 5) and aged mice housed in ENR or STD for one year (Aged ENR; Aged STD; n = 7). **a**, Aging decreases global CpG methylation in STD but not in ENR mice (*p*-values from two-way ANOVA with Dunnett’s post hoc test; depicted are means and s.e.m.). **b**, Aging and ENR induce opposite DNA methylation changes. Overlap of aging-induced hypoCpG with ENR-induced hyperCpG in aged mice (top left) and aging-induced hyperCpGs with ENR-induced hypoCpGs (top right). Overlap of aging- and ENR-induced CpH methylation changes (bottom). **c**, Age- and ENR-dependent changes of the distributions of methylation percentages of the 13,314 CpGs and 3,196 CpHs counteracted by ENR. Medians of methylation percentages per group are highlighted as black dots (CpGs; Young STD: 71.05 %, Young ENR: 68.57 %, Aged STD: 44.03 %, Aged ENR: 67.93 %. CpHs; Young-STD: 0 %, Young-ENR: 0 %, Aged-STD: 32.85 %, Aged ENR: 0 %). **d**, Heatmaps depicting the absolute methylation percentages in the individual groups at individual CpGs and CpHs. While aged STD mice show distinct methylation at individual sites, aged ENR mice show methylation levels similar to young animals.

To analyze whether ENR counteracts aging-induced DNA methylation changes at specific genomic loci, we determined cytosines where the methylation change induced by life-long ENR (difference between aged ENR and aged STD mice) was opposite to the effect of aging (difference between aged STD and young STD mice). From all CpGs that were hypomethylated with aging, 31.60 % were significantly hypermethylated in aged ENR mice compared to aged STD mice (Fig. 2b). Similarly, from all CpGs hypermethylated with aging, 32.17 % were hypomethylated by ENR. In the CpH context, 62.86 % of aging-induced hypermethylated CpHs and 7.24 % of hypomethylated CpHs were changed by ENR in the opposite direction than by aging (Fig. 2b). In total, 31.73 % of all age-related dmCpGs and 48.42 % of all aging-induced dmCpHs were counteracted by ENR. Among those cytosines, the vast majority (85.81 % of CpGs; 98.47 % of CpHs) did not show differences between young and aged ENR mice, confirming that, in ENR mice, methylation at these sites does not change with aging.

To compare magnitudes of methylation changes induced by aging and ENR, the absolute methylation percentages of the 13,314 dmCpGs and 3,196 dmCpHs at which ENR counteracted aging effects were plotted (Fig. 2c–d). Compared to the young animals, the aged STD mice exhibited a loss of highly methylated CpGs, which resulted in an accumulation of low or unmethylated CpGs and 27 % lower median CpG methylation levels. In contrast, the animals housed in ENR for one year showed distributions and median CpG methylation percentages similar to young animals. Likewise, aged STD mice possessed different distributions and a 33 % increase in the median CpH methylation percentage compared to young animals (Fig. 2d-e), while aged ENR mice were similar to young animals. These results suggest that ENR restores methylation at those age-sensitive CpGs and CpHs to the levels observed in young animals.

To increase temporal resolution of age-related DNA methylation changes, we integrated the data from the independent first experiment with 4.5-month-old (hereafter referred to as middle-aged) mice housed in ENR or STD for three months (compare Fig. 1). The CpG and CpH methylation levels of middle-aged mice were similar to young animals and did not show the age-related loss of CpG methylation observed in aged STD mice (Supplementary Fig. 4a). Further comparison of the effects of life-long ENR with age-related changes that occured from middle-aged STD to aged STD mice showed that the pattern of dmCpGs and dmCpHs that were counteracted by ENR was similar to that observed for young animals (Supplementary Fig. 4b). Additionally, 43.71 % of CpGs and 56.32 % of CpHs at which ENR counteracted aging were also differentially methylated between middle-aged STD mice and aged STD mice. This suggests that age-related epigenetic changes became pronounced only after an age of 4.5 months and that the environmental influence on age-related methylation changes is itself age-dependent.

### Genomic distribution of environmentally sensitive age-related DNA methylation changes

To further characterize the influence of ENR on aging-induced epigenetic reconfigurations, we analyzed the genomic distribution of CpGs and CpHs at which ENR counteracted age-related methylation changes. Both dmCpGs and dmCpHs were distributed over all chromosomes in the genome (Fig. 3a), with dmCpGs depleted at exons, CpG islands, CpG island shores, promoters and super-enhancers, but significantly enriched at introns, inter-genic regions and enhancers (Fig. 3b). The distribution of dmCpHs over genomic features was similar to dmCpGs, but they were not enriched at enhancers. Because DNA methylation of enhancers is particularly involved in the regulation of gene expression by interaction with transcription factors^29^, we performed a transcription factor motif analysis of the 1,472 dmCpGs located within enhancers. The only significantly enriched transcription factor was Mecp2 (Fig. 3c), which binds methylated cytosines via its methyl-CpG-binding domain^30^. Mecp2 was also the strongest enriched transcription factor when motif enrichment analysis was performed on all 13,314 dmCpGs (independent of genomic location), which suggested that Mecp2 binding occurs genome-wide and is not restricted to dmCpGs located within enhancers (Supplementary Data 5). In contrast to dmCpGs, no enrichment of Mecp2 motifs was found at dmCpHs (Fig. 3c).

**Fig. 3:**
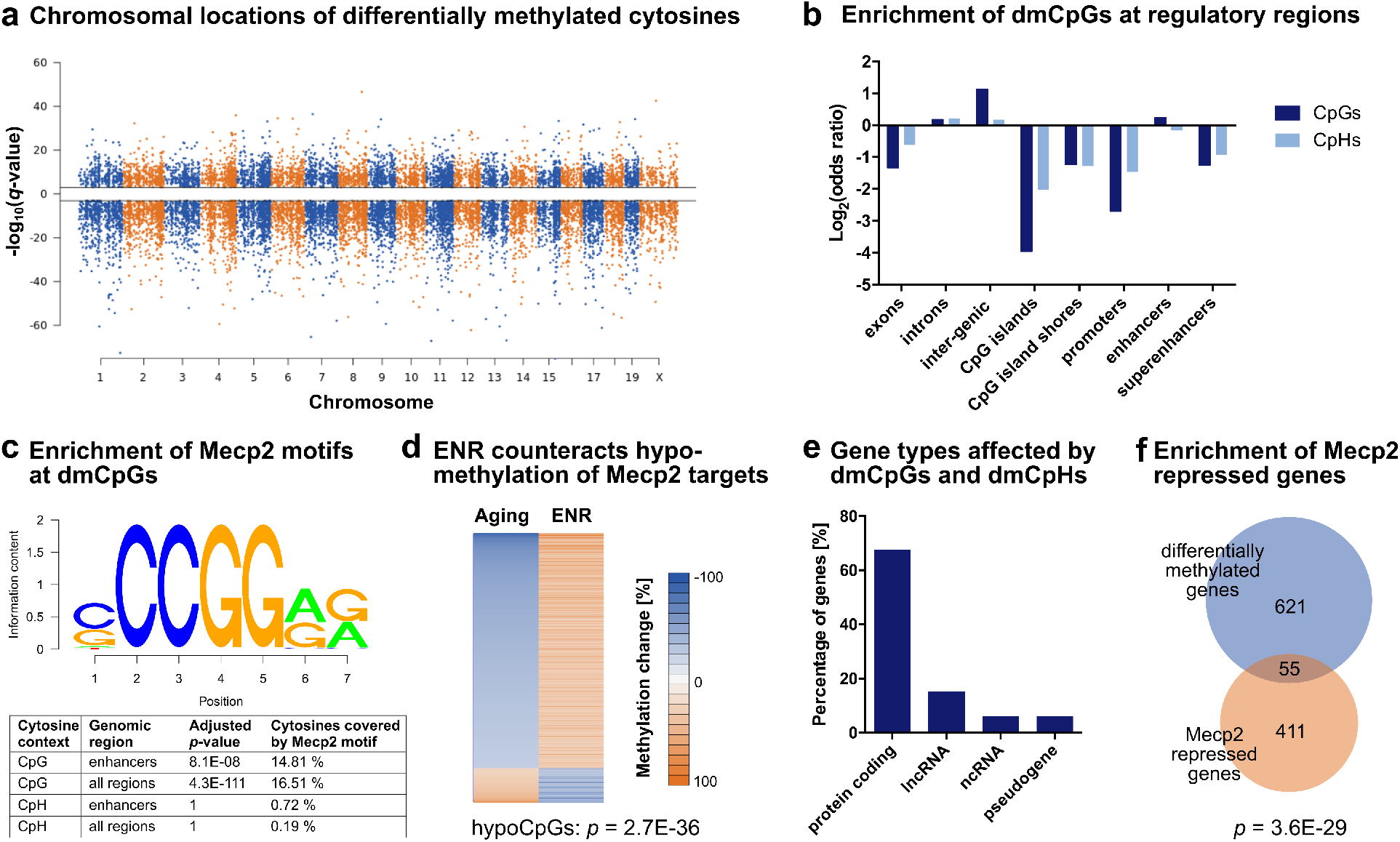
Genomic distribution of cytosines at which ENR counteracts age-related methylation changes. **a**, Manhattan plot depicting the distribution of dmCpGs and dmCpHs over chromosomes. *Q*-values indicate the significance of the aging-induced methylation changes with the sign corresponding to the direction of the change (negative -log_10_(*q*-value) for hypomethylated cytosines; positive -log_10_(*q*-value) for hypermethylated cytosines). **b,** CpGs and CpHs are depleted at CpG islands, CpG island shores, promoters and exons (log_2_(odds ratio) < 0), but enriched at introns and inter-genic regions of the genome (log_2_(odds ratio) > 0). Additionally, CpGs but not CpHs are enriched at enhancer regions. Adjusted *p* < 0.001 for all regions, except CpHs at enhancers (adjusted *p* = 0.061). **c,** Transcription factor binding analysis revealed enrichment of Mecp2 motifs at dmCpGs located within enhancers (1,472 CpGs) as well as at all regulated 13,314 dmCpGs (independent of genomic location). Mecp2 motifs were not enriched at dmCpHs. Depicted are the Mecp2 motif as annotated in the database motifDb and the results of the enrichment analysis. **d,** The 2,198 dmCpGs that fall into Mecp2 targets sites are enriched with CpGs that were hypomethylated with aging (*p*-value from hypergeometric test compared to all 13,314 CpGs). **e,** Classification of gene targets of cytosines at which ENR counteracts aging-induced methylation changes (676 differentially methylated genes). **f,** Previously identified Mecp2-repressed genes^32^ are enriched among the differentially methylated genes (*p*-value from hypergeometric test).

To validate the enrichment of Mecp2 binding sites at dmCpGs, we used two existing Mecp2 ChIP-sequencing datasets derived from adult mouse cortex and prefrontal cortex, respectively^31^. Similar to Mecp2 motifs, ChIP-derived Mecp2-bound genomic regions were significantly enriched at dmCpGs (hypergeometric tests; cortex: *p* < 1E-300; prefrontal cortex: *p* = 1.4E-286).

Because Mecp2 is known to specifically bind methylated cytosines, we analyzed the direction of the methylation change of Mecp2 target dmCpGs with aging and ENR. The vast majority of Mecp2 targets (87.31 %) were hypomethylated with aging and hypermethylated by ENR (Fig. 3d). Aging-induced hypomethylated CpGs were significantly enriched among the Mecp2 targets (*p* = 2.7E-36). These results suggest that ENR prevents the age-associated CpG hypomethylation of Mecp2 binding sites and, thereby, facilitates Mecp2 binding in the aged dentate gyrus.

Annotation of dmCpGs and dmCpHs to their associated gene targets identified 676 genes at which ENR counteracted age-related methylation changes. Two thirds of these genes were protein-coding genes, but also lncRNA and a minor percentage of ncRNAs (such as microRNAs) and pseudogenes changed DNA methylation (Fig. 3e). The differentially methylated genes were significantly enriched with genes that are known to be transcriptionally repressed by Mecp2^31^ (Fig. 3f), further supporting that ENR counteracts age-related DNA methylation changes at Mecp2 targets.

### Experience protects neuronal plasticity and neurogenesis-related genes from age-related DNA methylation changes

To identify aspects of brain aging that are sensitive to environmental stimulation, we performed GO and pathway enrichment analyses of genes at which ENR counteracted aging-induced methylation changes and generated an integrated map based on gene overlaps between enriched terms^33^. We identified three main clusters which were related to neuronal plasticity, cell communication and developmental programs (Fig. 4a; Supplementary Data 6). Among those, the “neuron plasticity” cluster contained the highest number of enriched terms. It comprised pathways related to structural plasticity of neurons, including axon guidance and synapse organization, as well as processes involved in neuronal activation, such as synaptic signaling and regulation of potassium channels. In the “cell communication” cluster, we found signaling pathways that involve secreted molecules, such as insulin and growth factors, and a group of enriched terms related to cell adhesion, including formation of cellular junctions by cadherins, adhesion to extracellular matrix and extracellular matrix remodeling by regulation of collagen formation. The third cluster contained pathways involved in organ morphogenesis but also pathways related to nervous system development, including neuronal fate specification and precursor proliferation.

**Fig. 4:**
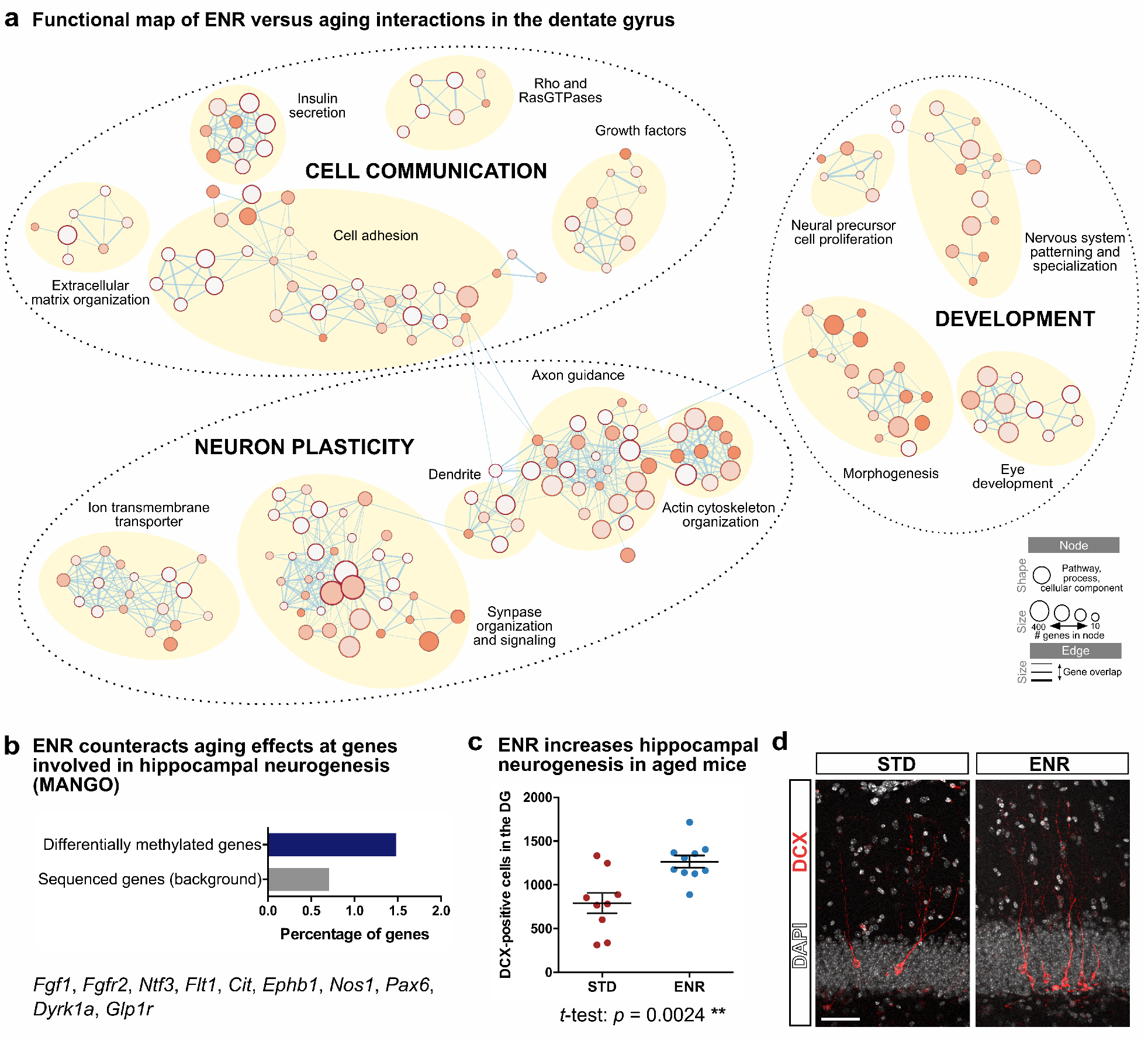
Functional enrichment connects environmentally sensitive age-related DNA methylation changes with neuronal plasticity and adult hippocampal neurogenesis. **a**, Integrated map of GO terms, SynGO terms, Reactome pathways and “WikiPathway” enriched among the 676 ENR-induced differentially methylated genes that counteract aging effects. Three main functional clusters were identified (dashed circles), each containing several groups of highly connected terms (yellow circles). Legend indicates node and edge sizes. **b**, Genes functionally involved in hippocampal neurogenesis are enriched among differentially methylated genes at which ENR counteracts age effects. Hypergeometric test: *p* = 0.023. **c**, Aged mice housed in ENR for one year have significantly more new-born, doublecortin (DCX)-expressing neurons in their dentate gyrus (DG). Depicted are individual sample points with mean and s.e.m. **d**, Representative image of fluorescent staining to detect new-born neurons in STD and ENR mice. Scale bar: 50 µm.

To correlate differential cytosine methylation with cellular changes in the brain, we investigated adult hippocampal neurogenesis which has a role in cognitive flexibility and decreases with aging^34^. We found that genes at which ENR counteracted age-related methylation changes were significantly enriched for genes functionally involved in adult hippocampal neurogenesis as annotated in MANGO (Fig. 4b). Accordingly, mice housed in ENR for one year showed a 60 % increase in the number of new-born neurons in the dentate gyrus compared to aged STD mice (Fig. 4c–d), suggesting that continuous ENR increases hippocampal neurogenesis throughout the lifespan. In contrast, only trends towards increased precursor proliferation and no change in total precursor cell numbers were observed in aged ENR compared to aged STD mice (Supplementary Fig. 5). This suggests that life-long ENR stimulates new-born neurons during an immature phase but has only minor effects on neural precursor cells, which is in accordance with what has been observed after shorter periods of ENR in young adult and aged mice^34^.

To analyze whether ENR protects from aging effects by actively changing DNA methylation throughout the lifespan, we overlapped genes where ENR counteracted DNA methylation changes in aged mice with DNA methylation changes induced by shorter periods of ENR in non-aged mice. In total, 24.02 % of genes that changed DNA methylation after four days of ENR in young mice and 30.03 % of genes that changed DNA methylation after three months of ENR in middle-aged mice overlapped with genes at which ENR counteracted age-related changes (Supplementary Fig. 6a). Additionally, ENR-induced differentially methylated genes were significantly enriched with genes that changed transcription after acute neuronal activation in the dentate gyrus^35^ (Supplementary Fig. 6b). These results might suggest that active ENR-induced DNA methylation changes at neuronal activity regulated genes counteract the development of age-related DNA methylation changes.

### ENR changes DNA methylation in mice at genes associated with age-related cognitive decline in the human brain

To investigate whether the genes at which ENR counteracted age-related DNA methylation changes in the mouse brain are also dysregulated in the human brain during aging and associated cognitive decline, we overlapped them with three previously published datasets derived from human prefrontal cortex tissue. Dataset 1 contained genes with DNA methylation changes associated with Alzheimer’s disease pathology^20^; dataset 2 comprised genes with changes in RNA levels associated with age-related cognitive decline^36^ and dataset 3 contained genes with proteomic changes associated with individual cognitive trajectories during aging^37^. Genes of all three datasets were significantly enriched among the genes at which ENR counteracted age-related DNA methylation changes (Fig. 5a). In total, 256 of the ENR-induced differentially methylated genes detected in the aged mouse brain were also dysregulated in the aged human brain (Fig. 5b; Supplementary Data 7). Among those genes, the most significant enriched biological process from GO analysis was the term “neurogenesis” (adjusted *p* = 7.9E-08; 53 genes) of the higher order group “nervous system development” (adjusted *p* = 5.0E-10; 72 genes). This suggests that the age-related dysregulation of genes involved in neurogenesis is conserved between mouse and human. The overlap of genes regulated by ENR in mice and genes dysregulated in human brains highlights the intriguing possibility that environmental stimulation could prevent age-related gene dysregulation and associated cognitive decline also in human brains.

**Fig. 5:**
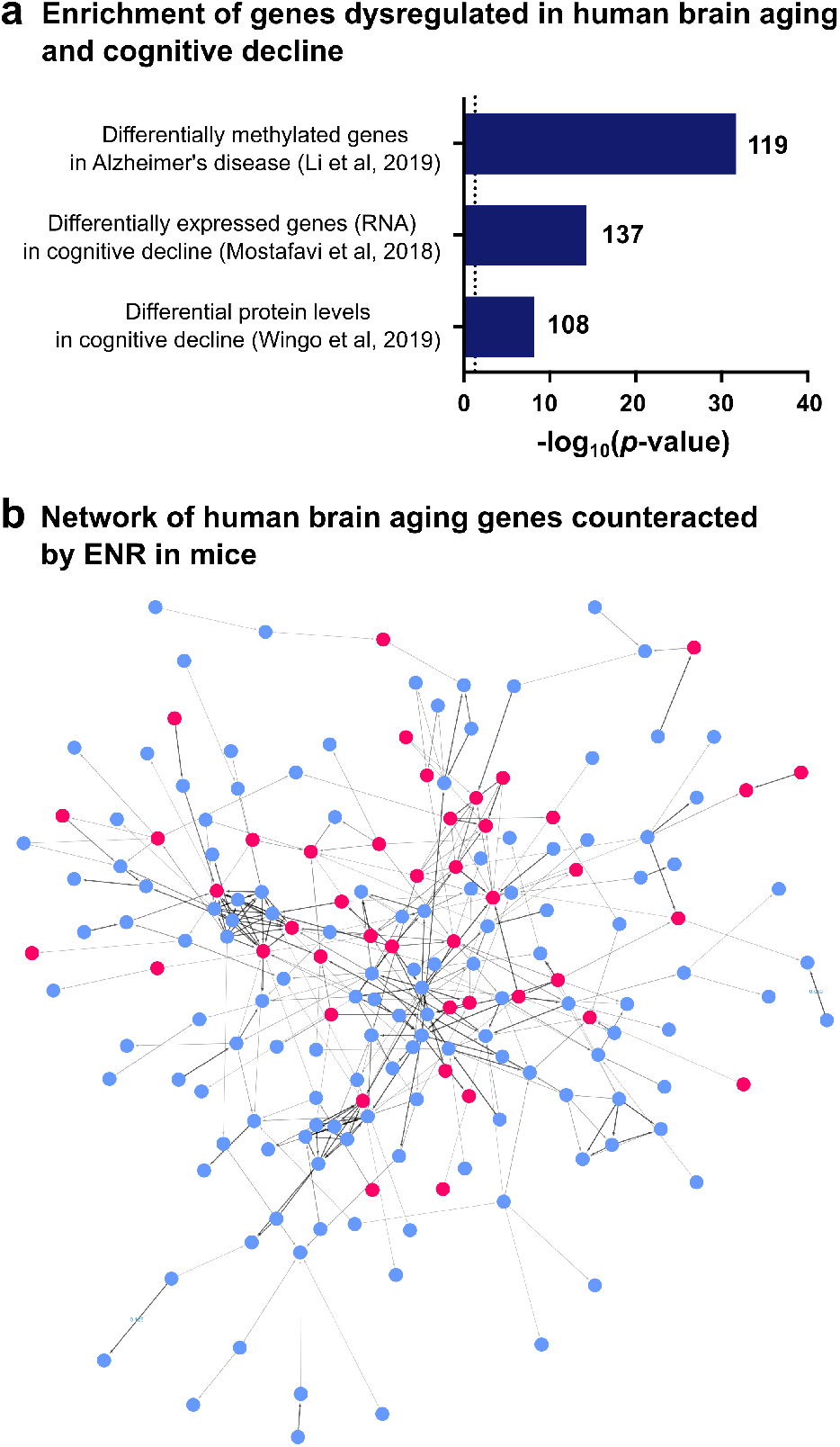
Genes at which ENR counteracts age-related methylation changes are dysregulated in the human brain in age-related cognitive decline. **a,** Genes with DNA methylation changes^20^, differential RNA^36^ or protein^37^ levels in human prefrontal cortex associated with age-related cognitive decline are significantly enriched among the genes that are differentially methylated with aging and ENR in the mouse dentate gyrus. Depicted are the -log_10_(*p*-values) from hypergeometric tests and the number of genes that overlapped between datasets. Dashed line indicates significance threshold of *p* = 0.05. **b,** STRING protein interaction network of human brain aging genes at which ENR counteracted age-related DNA methylation changes in the mouse brain. Genes involved in the highest enriched GO biological process “neurogenesis” are highlighted in red.

## Discussion

Aging-induced epigenetic changes are conserved among species^38^ and contribute to age-related brain dysfunctions, including reductions in cognition and hippocampal neurogenesis^9, 10^. Behavioral interventions and environmental stimulation are known to alter epigenetic modifications in the brain^15, 32, 39^, but their influence on age-related epigenetic changes was unclear. Here, we showed that lifelong ENR can counteract age-related DNA methylation changes in the hippocampal dentate gyrus. Many of the genes for which ENR prevented aging-induced differential methylation in the mouse brain were also known to be dysregulated in human brains with aging and related cognitive decline. Our results lend mechanistic support to the potential of behavioral interventions and lifestyle factors to prevent age-related epigenetic changes in the brain in order to improve brain health in old age.

We found that ENR prevented the aging-induced loss of CpG methylation genome-wide at binding sites of Mecp2. Since Mecp2 binds methylated cytosines with high affinity broadly across the genome^40–42^, these results suggested that aging is associated with reduced Mecp2 binding to neuronal genomes and that ENR enhances Mecp2 binding in aged brains. Mutations in Mecp2 cause Rett syndrome, which is characterized by severe encephalopathy and reduced lifespan^43^. Mecp2 is known to be abundant in neurons and crucial for neuronal development during embryogenesis and in the adult hippocampus^44, 45^, but a role of Mecp2 in brain aging had not yet been reported. At the molecular level, Mecp2 has been associated with transcriptional repression, locus-specific gene activation and regulation of alternative splicing^41, 42, 46^. Therefore, aging-induced loss of Mecp2 binding as a result of genome-wide CpG hypomethylation might be involved in the aging-associated transcriptional dysregulation and the aberrant splicing of neuronal genes that has been reported for the aged hippocampus^47^. On the other hand, a recent study suggested that Mecp2 is involved in the repression of endogenous repetitive elements in neuronal genomes^48^. ENR-induced methylation of Mecp2 targets could thus prevent the aberrant activation of repetitive elements which is known to contribute to genomic instability in aging^49^. Follow-up studies should experimentally confirm the interaction between CpG hypomethylation and Mecp2 binding during brain aging and identify molecular programs downstream of Mecp2 that are disrupted in the aged brain. Our results suggest not only a possible role of Mecp2 in age-related cognitive decline but also the potential of environmental stimulation to rescue Mecp2 binding in aged brains by DNA methylation of its target sites.

With the exception of a few regulated genes^50, 51^, the molecular mechanisms underlying ENR-stimulated brain plasticity were, until now, mostly unknown. Our functional enrichment analyses of differentially methylated genes in the aged mouse dentate gyrus have exposed several pathways potentially underlying the neuroprotective effects of ENR during aging. One such pathway was adult hippocampal neurogenesis and, since increasing neurogenesis promotes aspects of hippocampal function^52^, ENR-stimulated increases in hippocampal neurogenesis likely ameliorate age-related cognitive decline. Further investigation will be needed to determine the cell type specificity of ENR-induced DNA methylation changes in the dentate gyrus and their role in in hippocampal neurogenesis.

Since ENR increased adult hippocampal neurogenesis throughout the lifespan, differences in DNA methylation patterns between STD and ENR mice could reflect DNA methylation changes in mature neurons or ENR-induced differences in the cellular composition of the dentate gyrus. Based on our quantifications of new-born neuron numbers, supported by previous quantifications of total granule cell numbers in the dentate gyrus^53^, the percentage of new-born neurons among all neurons increased from 0.21 % in aged STD mice to 0.31 % in aged ENR mice. Because our DNA methylation analysis only considered cytosines with absolute DNA methylation differences larger than 25 % as dmCpGs or dmCpHs, DNA methylation differences as a result of increased percentage of new neurons in ENR mice would have not been detected. Second, because hippocampal neurogenesis sharply decreases in young adulthood and plateaus thereafter^54^, differences in DNA methylation patterns as a result of differences in newborn neuron numbers should have been visible in the 4.5-month-old ‘middle-aged’ mice. However, the middle-aged mice showed only minor age-related DNA methylation changes compared to young animals (Supplementary Fig. 4). Therefore, we conclude that the here described ENR-induced DNA methylation changes that counteracted age effects reflect DNA methylation changes in mature hippocampal neurons rather than changes in cellular composition of the dentate gyrus.

ENR combines physical and social incentives with continuous novelty and sensory stimulation^4^. Which aspects of ENR and the mechanisms how environmental stimulation prevents age-related DNA methylation changes remains to be investigated. Previous work has shown that exploration of novel environments leads to neuronal activation coupled with short-term and long-lasting changes in transcription and chromatin accessibility in activated hippocampal neurons^55, 56^. Additionally, isolated neuronal activation by electroconvulsive stimulation has been shown to lead to widespread changes in DNA methylation, RNA and open chromatin landscapes in the dentate gyrus^16, 35^. We showed that ENR-induced differentially methylated genes were significantly enriched for neuronal activity regulated genes, indicating that some of the ENR-induced DNA methylation changes are triggered by neuronal activation. Similarly, Penner et al. reported that age-related DNA methylation changes at the promoter of the neuronal activity-induced gene *Egr1* were reversed in the hippocampus by acute exploration of a novel environment^19^. We found that many of the genes at which ENR counteracted age-related DNA methylation changes were also differentially methylated by acute (four days) ENR in non-aged brains or by three months ENR in middle-aged mice. Therefore, the rejuvenating effects of ENR on age-related DNA methylation changes are potentially mediated by lifelong, repeated neuronal activation through continuous novelty stimulation in ENR. Future studies should specify how ENR interacts with age-related DNA methylation changes and address whether short-term ENR of aged animals is sufficient to epigenetically rejuvenate the hippocampus.

ENR has an enormous potential to prevent or counteract brain dysfunctions in aged animals, including synaptic plasticity, hippocampal neurogenesis and cognitive abilities^57–59^. Using genome-wide DNA methylation sequencing at single nucleotide resolution, we here demonstrated that ENR also restores a substantial number of age-related DNA methylation changes in the hippocampus. Our data show the rejuvenating effects of ENR at the epigenetic level and provide a potential mechanism how active interaction with the environment supports and promotes brain function throughout the lifespan.

## Methods

### Animals and environmental enrichment

Female C57BL/6JRj mice were ordered from Janvier Labs and maintained on a 12 h light/dark cycle at the animal facility of the Center for Regenerative Therapies Dresden. Food and water were provided *ad libitum*. At an age of 6 weeks, mice were randomly distributed to an enriched environment or control cages. The enriched environment was a 0.74 m^2^ enclosure equipped with tunnels and plastic toys. Toys were rearranged once per week, but not in the last four days before analysis. In every experiment, ten mice were housed together in an enriched environment at the same time. Control mice stayed in standard polycarbonate cages (Type II, Tecniplast) in groups of five mice per cage. Dirty bedding material and toys were replaced once per week. The entire enriched environment was cleaned once per month. All experiments were conducted in accordance with the applicable European and national regulations (Tierschutzgesetz) and were approved by the local authority (Landesdirektion Sachsen).

### Reduced Representation Bisulfite Sequencing (RRBS)

Genomic DNA was isolated from micro-dissected dentate gyrus tissue using the QIAamp DNA Micro Kit or the Allprep DNA/RNA Micro Kit (both QIAGEN) following the manufacturer’s manuals. RRBS libraries were prepared using the Premium RRBS Kit (Diagenode) and purified twice using Agencourt AMPure XP beads (Beckman Coulter; 1X bead volume). Quality and concentration of RRBS libraries were determined using the High Sensitivity NGS Fragment Analysis Kit (Advanced Analytical) and a fragment analyzer with capillary size of 33 cm (Advanced Analytical). Sequencing was performed using a HiSeq2500 (experiment with middle-aged animals; Fig. 1) or a Nextseq (experiment with young and aged animals; Fig. 2-5) platform in a 75 bp single end mode with a minimum sequencing depth of 15 million reads per sample.

### Bioinformatic data analysis

#### Calculation of differential DNA methylation

Fastq reads were trimmed using Trim Galore 0.4.4 and the function *Cutadapt* 1.8.1. To remove cytosines that were filled in during end preparation, an additional 2 bp were cut off from every sequence with detected adapter. Trimmed reads were mapped against mm10 using Bismark 0.19.0^60^.

Global DNA methylation levels are the means of the methylation percentages over all cytosines that were covered by RRBS with at least 10 reads in all samples of the respective experiments. Separation into the cytosine contexts CpG and CpH (CHH and CHG) was performed based on context annotations extracted from Bismark.

Detection of differentially methylated cytosines was performed using methylKit 1.5.2^61^. Briefly, methylation levels were extracted from sorted Binary Alignment Map files using the function *processBismarkAln*. Data was filtered for CpGs with a minimum coverage of ten reads and a maximum coverage of 99.9% percentile using the function *filterByCoverage*. Using the function *unite*, CpGs were selected that were sufficiently covered in at least three samples per group. Differentially methylated cytosines were identified using the *methDiff* function applying the chi-squared test, a significance threshold of *q* < 0.001 and a threshold for absolute DNA methylation differences higher than 25 %.

Differentially methylated cytosines were annotated to the gene with the nearest transcription start site using data tables downloaded from Ensembl BioMart (as of 01.05.2019)^62^. Genes with at least four annotated differentially methylated cytosines were considered differentially methylated genes. Gene names used in this study are Ensembl gene names. Corresponding Entrez identifiers were retrieved from the Bioconductor package AnnotationDbi^63^ using Ensembl gene identifiers as keys. Mapping of mouse genes to homologous human genes was performed using data tables downloaded from Ensembl BioMart^62^.

#### Genomic feature annotation

Genomic coordinates of CpG islands, exons and introns were downloaded from the UCSC genome browser^64^. CpG island shores were defined as 2 kb upstream and downstream of a CpG island. Promoter regions were determined as ± 1 kb from the transcription start sites of all known transcripts. Enhancer and transcription factor binding sites were downloaded from the Ensembl Regulatory Build track of the UCSC genome browser^65^. Superenhancer regions are from adult mouse cortex and were retrieved from dbSUPER^66^. Overlaps of cytosines with genomic regions or other cytosines were performed using the functions *subsetByOverlaps* or *findOverlaps* of the R package Genomic Ranges 3.7^67^. Genomic feature enrichment was analyzed using count tables of feature coverage of differentially methylated cytosines and all cytosines covered by RRBS (background). Odds ratios and *p*-values of feature enrichment were calculated by fitting a general linear model with binomial distribution using the function *glm* in R. Odds ratios are the exponential coefficients of the model.

#### Transcription factor binding analysis

The number of differentially methylated cytosines that overlap with position weight matrices of transcription factor binding motifs was determined using the *Biostrings* package with a minimum match score of 90 %. Transcription factor motifs were retrieved from motifDb^68^. Motif enrichment for each transcription factor was tested by applying hypergeometric tests using the function *phyper(q, m, n, k) + dhyper(q, m, n, k)* with q = number of differentially methylated cytosines overlapping with the motif, m = number of background cytosines overlapping with the motif, n = number of background cytosines not overlapping with the motif and k = total number of differentially methylated cytosines. Multiple testing correction of *p*-values was performed using the FDR method.

For conformation of Mecp2 binding sites, Mecp2 ChIP-Seq data were downloaded from GEO (GSE67293) and overlapped with dmCpGs using Genomic Ranges 3.7. Hypergeometric test for the enrichment analysis was performed as described for the motif analysis.

### Functional gene enrichment analyses

Enrichment for gene ontology (GO) cellular components was analyzed using the online tool GREAT^69^. Pathway analysis was performed using the R package ReactomePA^70^ with a minimum of 5 and a maximum of 300 genes per pathway. Enrichment for genes involved in synaptic processes was performed using the SynGO online tool (https://www.syngoportal.org/) with default settings. All enrichment analyses were performed with differentially methylated genes as query lists and all genes covered by RRBS as background lists.

Gene set enrichment was analyzed by performing hypergeometric tests using the function *phyper(q, m, n, k) + dhyper(q, m, n, k)* in R with q = number of differentially methylated genes overlapping with the gene set, m = number of background genes overlapping with the gene set, n = number of background genes not overlapping with the gene set and k = total number of differentially methylated genes. Genes involved in adult hippocampal neurogenesis were downloaded from the Mammalian Adult Neurogenesis Gene Ontology (MANGO) v3.2^14^. The gene list was filtered for genes with reported positive or negative effect on neurogenesis after protein manipulation or gene manipulation (in total 283 genes). Genes or cytosines with known aging-induced DNA methylation changes were extracted from^38, 71, 72^ for peripheral tissues and from ^18, 28^ for the hippocampus.

The enrichment map was generated according to Merico et al.^33^ using genes that contain at least four differentially methylated cytosines (dmCpGs or dmCpHs). GO biological process, GO cellular component, Reactome, and Wikipathways enrichment analyses were performed in g:Profiler (https://biit.cs.ut.ee/gprofiler) using gene ensembl identifiers. The results from SynGO and MANGO enrichment analysis were integrated into the gmt file downloaded from g:Profiler. The network was generated using the app Enrichment Map 3.2.0 in cytoscape 3.7.1 using a *q*-value threshold (FDR) of 0.05. Clusters were manually annotated. Groups with fewer than six connected terms were not drawn. The STRING protein interaction network was generated inside cyotscape 3.7.1 using human ensembl identifiers with the plugin STRING (protein query) at a confidence score cutoff of 0.5 with automated enrichment analysis.

#### Data and code availability

Sequencing data are deposited at GEO (accession number pending). Sequencing data were analyzed using in-house R scripts which are available upon request.

### Analysis of adult hippocampal neurogenesis

Tissue fixation and immunofluorescent stainings for the detection of new-born neurons were performed as previously described^34^. Briefly, mice were anesthetized with 100 mg/kg ketamine (WDT) and 10 mg/kg xylazin (Serumwerk Bernburg AG) and transcardially perfused with 0.9 % sodium chloride. Brains were removed from the skull and one hemisphere fixed in 4 % paraformaldehyde prepared in phosphate buffer (pH 7.4) overnight at 4°C. Brains were incubated in 30 % sucrose in phosphate buffer for two days and cut into 40 µm coronal sections using a dry-ice-cooled copper block on a sliding microtome (Leica, SM2000R). Sections were stored at 4 °C in cryoprotectant solution (25 % ethyleneglycol, 25 % glycerol in 0.1 M phosphate buffer, pH 7.4). For immunofluorescent stainings, sections were washed and unspecific binding sites were blocked in phosphate buffered saline supplemented with 10 % donkey serum (Jackson Immuno Research Labs) and 0.2 % Triton X-100 (Carl Roth) for 1 h at room temperature. Primary antibodies were applied overnight at 4 °C as follows: polyclonal goat anti-doublecortin (1:250; Santa Cruz), polyclonal rat anit-Ki67 (1:500; eBioscience), polyclonal goat anti-Sox2 (1:500; Santa Cruz). The secondary antibodies anti-goat Cy3 and anti-rat 488 (1:1000; Jackson Immuno Research Labs) were incubated for 2 h at room temperature. Antibodies were diluted in phosphate buffered saline supplemented with 3 % donkey serum and 0.2 % Triton X-100. Nuclei were labeled with 4’,6-diamidino-2-phenylindole (DAPI; 1:4000; Jackson Immuno Research Labs). Sections were mounted onto glass slides and cover-slipped using Aquamount (Polysciences Inc.). Stainings were imaged using a Zeiss Apotome equipped with an AxioCam MRm camera and the software AxioVision 4.8 (Zeiss). Cells were quantified in every sixth section along the rostro-caudal axis of the dentate gyrus.

### Statistics

The statistical tests used are reported in the specific results or methods section. All *t*-tests were two-sided and performed using GraphPad Prism 6.0. When performing multiple comparisons, *p*-values were adjusted using FDR corrections. All measurements were taken from distinct samples.

## Acknowledgments

This study was financed from basic institutional funds. S.Z. was supported by an EMBO Short-Term Fellowship and a fellowship from the International Max Planck Research School on the Life Course, Berlin. The authors thank Hongjun Song for discussion and methodological support.

## Author contributions

Conceptualization – S.Z., G.K.; Investigation/ Visualization – S.Z.; Methodology/ Formal analysis – S.Z., R.W.O; Software/ Data Curation – R.W.O., M.L., S.Z.; Resources – A.D., G.K.; Writing original draft – S.Z.; Writing review and editing – S.Z., R.W.O., G.K..

## Supplementary Information

**Supplementary Fig. 1.**
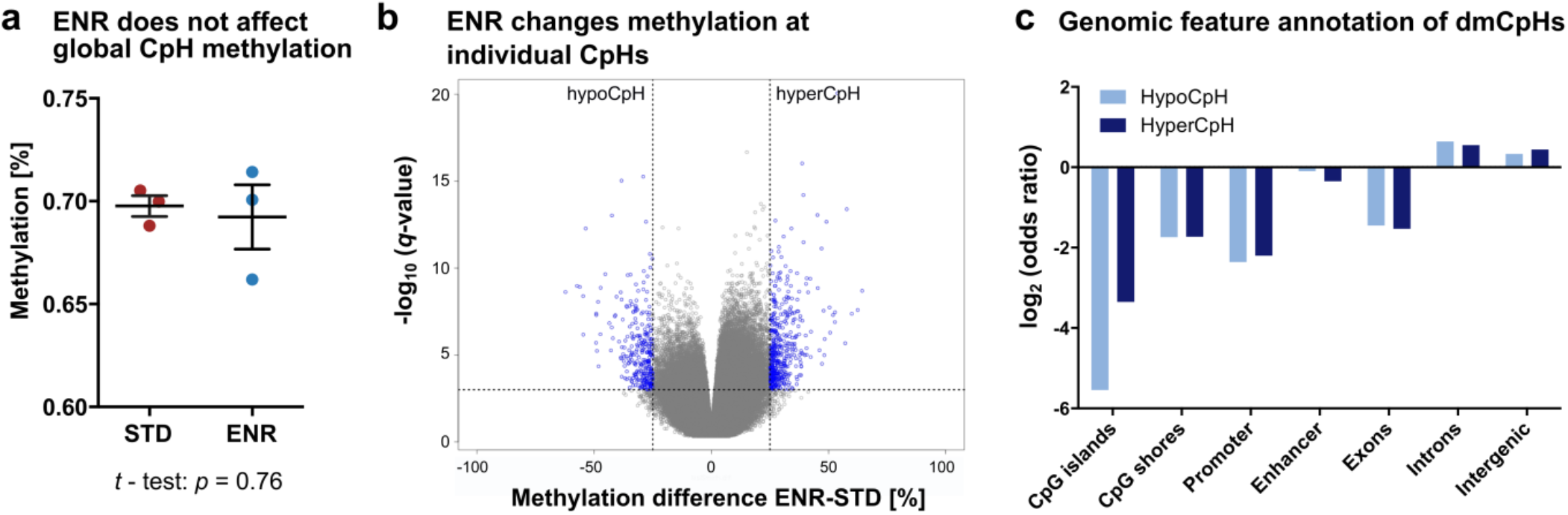
Environmental enrichment elicits locus-specific CpH methylation changes in the adult, non-aged dentate gyrus. DNA methylation profiles from 4.5 months old mice housed in ENR or STD for three months. **a**, ENR does not change global CpH methylation. Depicted are individual data points with mean and s.e.m. **b**, Volcano plot depicting differentially methylated CpHs (dmCpHs) in blue. In total, 750 dmCpHs were detected (0.019 % of covered CpHs). Of those, 38.93 % were hypomethylated (hypoCpH) and 61.07 % were hypermethylated (hyperCpH) in ENR mice. **c**, Genomic distribution of hypoCpH and hyperCpH (adjusted *p* < 0.001 for CpG islands, CpG island shores, promoters, exons, introns; adjusted *p* < 0.05 for intergenic regions; enhancer: adjusted *p* = 0.65 for hypoCpH, adjusted *p* = 0.26 for hyperCpH).

**Supplementary Fig. 2.**
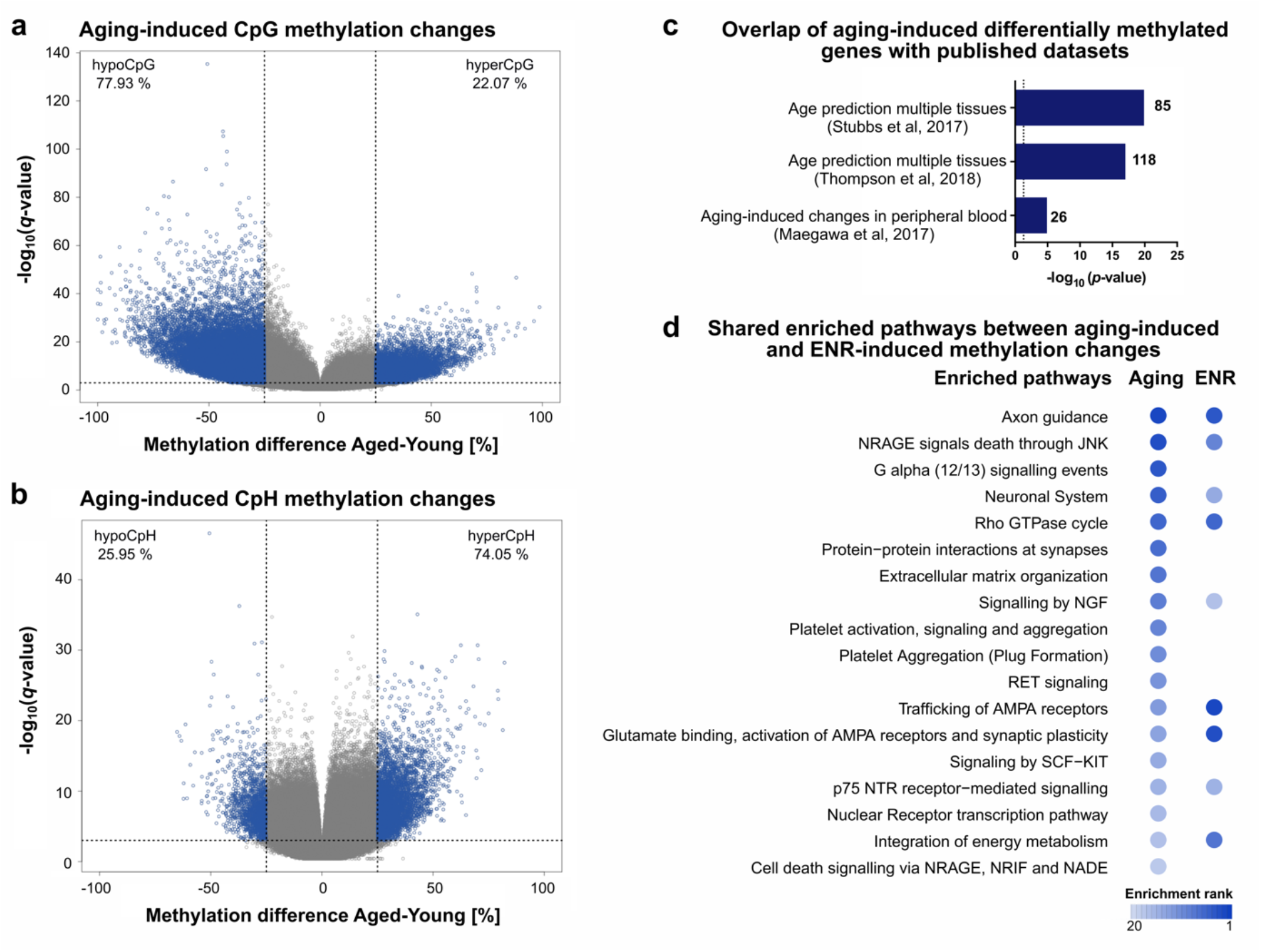
Age-related CpG and CpH methylation changes in the dentate gyrus. **a**, Volcano plot depicting CpGs where aging changed methylation in STD mice in blue (n = 5 for young mice; n = 7 for aged mice). In total, 5.51 % of all covered CpGs were dmCpGs. Age-related CpG hypomethylation (77.93 % of dmCpGs) is more prominent than CpG hypermethylation (22.07 % of dmCpGs). **b**, Aging changed methylation at 0.17 % of CpHs in the genome. The vast majority of dmCpHs were hypermethylated in the aged dentate gyrus (74.05 %). **c**, Enrichment of age-related differentially methylated genes (CpG and CpH context) from published dataset among genes with age-related CpG or CpH methylation changes in the dentate gyrus (*p*-values from hypergeometric tests). **d**, Top 18 pathways from Reactome pathway analysis with age-related differentially methylated genes (in total 3983 genes). Pathways were ordered by significance of the enrichment and colored based on the rank of significance. Among the top 18 enriched pathways from age-related methylation changes are also the highest enriched pathways from ENR-induced DNA methylation changes in non-aged mice after three months of ENR.

**Supplementary Fig. 3.**
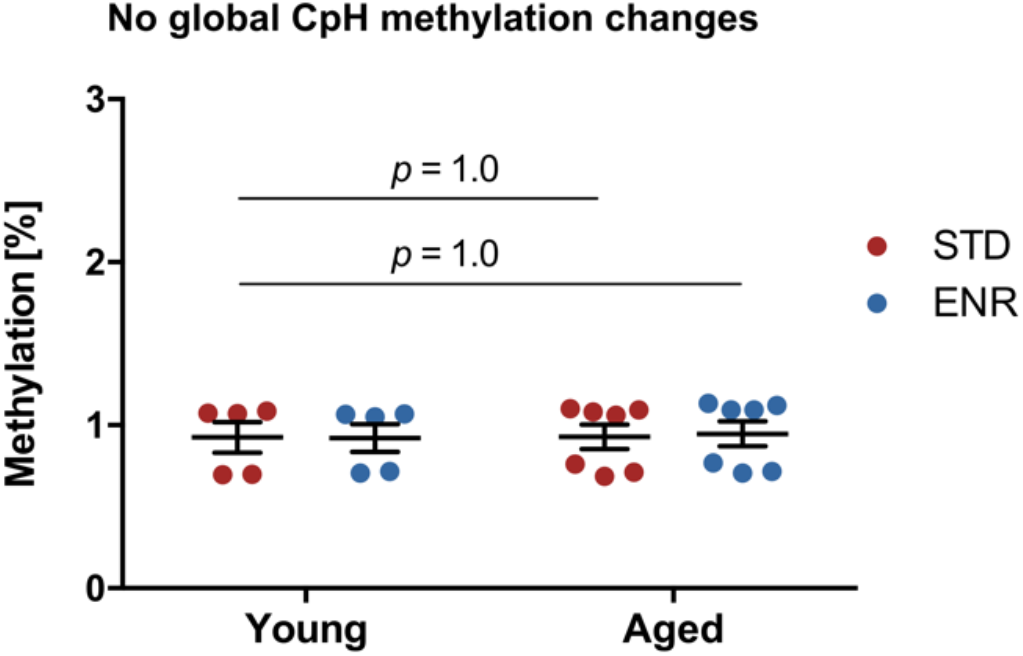
No interaction between environmental enrichment and aging in global CpH methylation. Aging and ENR do not change global CpH methylation (mean over 246,878 CpHs). The *p*-values are from two-way ANOVA with Dunnett’s post hoc test.

**Supplementary Fig. 4.**
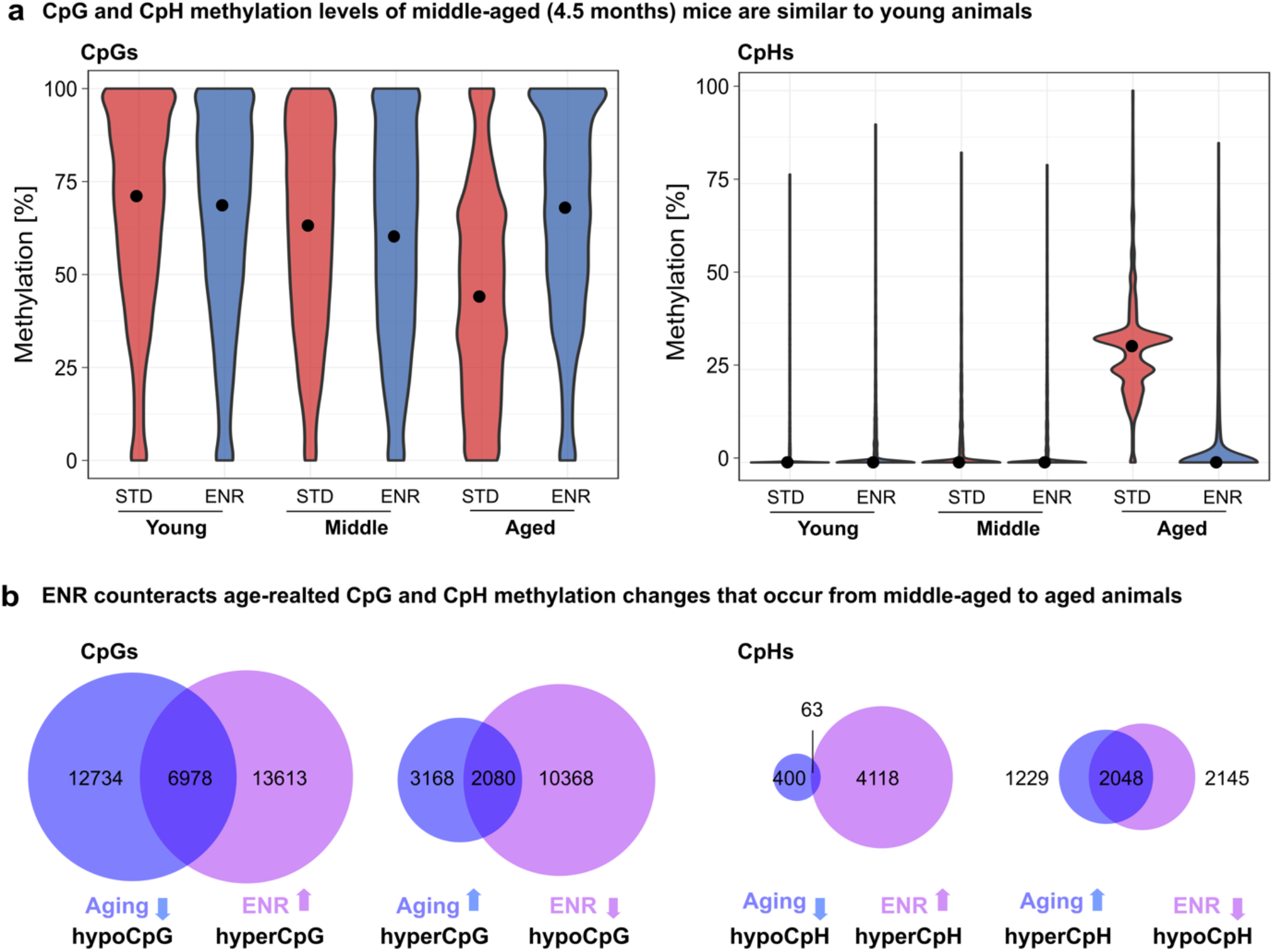
The interaction of age-related and ENR-induced DNA methylation changes between middle-aged and aged animals. **a**, STD and ENR housed middle-aged animals (4.5 months) show CpG and CpH methylation levels similar to young animals. **b**, In total, 36.29 % of CpG methylation differences between 4.5-month-old and 14-month-old STD mice and 56.44 % of CpH methylation changes were counteracted by ENR.

**Supplementary Fig. 5.**
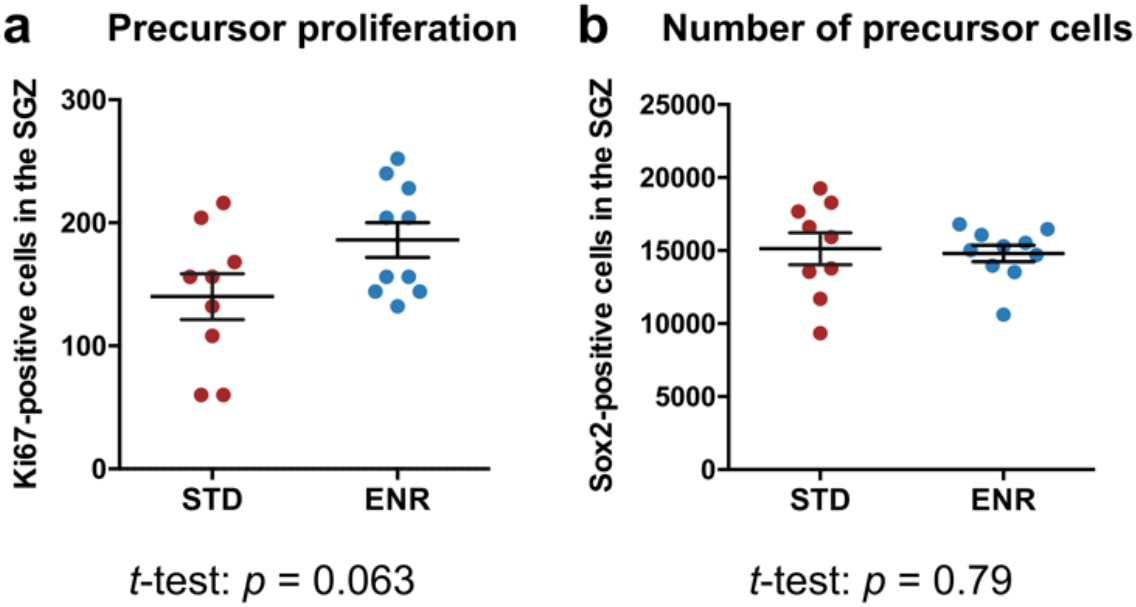
Influence of life-long ENR on neuronal precursor cells in the dentate gyrus. **a**, Quantification of the numbers of proliferating, Ki67-positive precursor cells in the subgranular zone (SGZ). **b**, No difference in total numbers of Sox2-positive precursor cells in the SGZ between aged mice housed in STD or ENR for one year.

**Supplementary Fig. 6.**
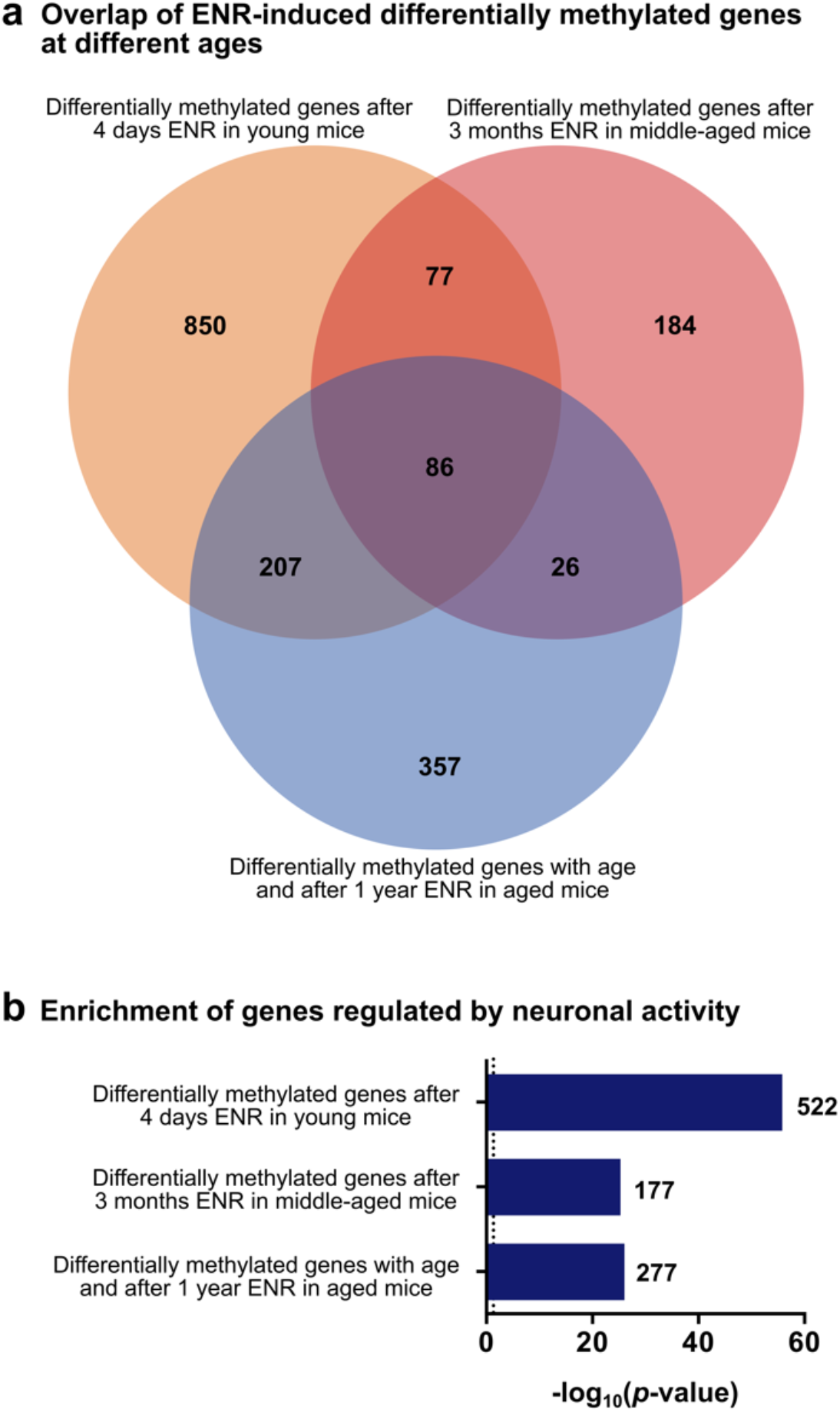
Genes at which ENR counteracts age-related DNA methylation changes overlap with ENR-induced methylation changes in the non-aged brain and with neuronal activity regulated genes. **a**, Venn diagram with number of overlapping genes between groups. **b**, Enrichment of genes that change RNA levels after activation of neurons in the dentate gyrus by electroconvulsive stimulation. Depicted are the -log_10_(*p*-values) and the number of overlapping genes for every dataset.

## Captions for Supplementary Data files

**Supplementary Data 1**

Differentially methylated cytosines (sheet 1) and differentially methylated genes (sheet 2) in the dentate gyrus of 4.5-month-old mice after three months of ENR housing. Data related to Fig. 1.

**Supplementary Data 2**

Results of GO and pathway enrichment analyses with gene targets of ENR-induced dmCpGs in 4.5 months old mice. Listed are significantly enriched pathways of “Reactome” pathway enrichment analysis (sheet 1), significantly enriched GO terms from GO cellular component enrichment analysis (sheet 2), significantly enriched GO terms of SynGO biological process enrichment analysis (sheet 3) and differentially methylated genes with functional annotation in MANGO (sheet 4) and all 373 differentially methylated genes (sheet 5). Data related to Fig. 1.

**Supplementary Data 3**

Lists of age-related differentially methylated cytosines in STD mice (sheet 1) and ENR-induced differentially methylated cytosines in aged mice (sheet 2). Data related to Fig. 2 and Supplementary Fig. 1.

**Supplementary Data 4**

Significantly enriched pathways of age-related differentially methylated genes from “Reactome” pathway enrichment analysis. Data related to Supplementary Fig. 2.

**Supplementary Data 5**

Results of transcription factor motif enrichment analysis of age-related dmCpGs counteracted by ENR. Listed are significantly enriched transcription factor motifs in all 13,314 dmCpGs (sheet 1), enriched motifs in dmCpGs located within enhancers (sheet 2) and locations and methylation changes of Mecp2 target dmCpGs (sheet 3). Data related to Fig. 3.

**Supplementary Data 6**

Results of enrichment analysis used for generation of the enrichment map (sheet 1) and list of the 676 genes at which ENR counteracted age-related DNA methylation changes (sheet 2). Data related to Fig. 4.

**Supplementary Data 7**

List of genes that are dysregulated with age-related cognitive decline in humans and regulated by ENR in mice (sheet 1) and results of STRING enrichment analysis (sheet 2). Data related to Fig. 5.

## References

1. Phillips, C. Lifestyle Modulators of Neuroplasticity: How Physical Activity, Mental Engagement, and Diet Promote Cognitive Health during Aging. Neural Plast. (2017).

2. Cabeza, R. et al. Maintenance, reserve and compensation: the cognitive neuroscience of healthy ageing. Nat. Rev. Neurosci. 19, 701–710 (2018).

3. Stern, Y. et al. Whitepaper: Defining and investigating cognitive reserve, brain reserve, and brain maintenance. Alzheimer’s Dement. 1–7 (2018).

4. Kempermann, G. Environmental enrichment, new neurons and the neurobiology of individuality. Nat. Rev. Neurosci. (2019).

5. Rogers, J., Renoir, T. & Hannan, A. J. Gene-environment interactions informing therapeutic approaches to cognitive and affective disorders. Neuropharmacology 145, 37– 48 (2019).

6. Fan, X., Wheatley, E. G. & Villeda, S. A. Mechanisms of Hippocampal Aging and the Potential for Rejuvenation. Annu. Rev. Neurosci. 40, 251–272 (2017).

7. Blasco, M. A., Partridge, L., Serrano, M., Kroemer, G. & Lo, C. The Hallmarks of Aging. Cell 153, 1194–1217 (2013).

8. Horvath, S. DNA methylation age of human tissues and cell types. Genome Biol. 14, R115 (2013).

9. Oliveira, A. M. M., Hemstedt, T. J. & Bading, H. Rescue of aging-associated decline in Dnmt3a2 expression restores cognitive abilities. Nat. Neurosci. 15, 1111–1113 (2012).

10. Gontier, G. et al. Tet2 Rescues Age-Related Regenerative Decline and Enhances Cognitive Function in the Adult Mouse Brain. Cell Rep. 22, 2094–2106 (2018).

11. Kaas, G. A. et al. TET1 Controls CNS 5-Methylcytosine Hydroxylation, Active DNA Demethylation, Gene Transcription, and Memory Formation. Neuron 79, 1086–1093 (2013).

12. Feng, J. et al. Dnmt1 and Dnmt3a maintain DNA methylation and regulate synaptic function in adult forebrain neurons. Nat. Neurosci. 13, 423–430 (2010).

13. Nelson, E. D., Kavalali, E. T. & Monteggia, L. M. Activity-Dependent Suppression of Miniature Neurotransmission through the Regulation of DNA Methylation. J. Neurosci. 28, 395–406 (2008).

14. Weaver, I. C. G. et al. Epigenetic programming by maternal behavior. Nat. Neurosci. 7, 847–54 (2004).

15. Halder, R. et al. DNA methylation changes in plasticity genes accompany the formation and maintenance of memory. Nat. Neurosci. 19, 102–110 (2016).

16. Guo, J. U. et al. Neuronal activity modifies the DNA methylation landscape in the adult brain. Nat. Neurosci. 14, 1345–1351 (2011).

17. Cholewa-Waclaw, J. et al. The Role of Epigenetic Mechanisms in the Regulation of Gene Expression in the Nervous System. J. Neurosci. 36, 11427–11434 (2016).

18. Masser, D. R. et al. Sexually divergent DNA methylation patterns with hippocampal aging. Aging Cell 16, 1342–1352 (2017).

19. Penner, M. R. et al. Age-related changes in Egr1 transcription and DNA methylation within the hippocampus. Hippocampus 26, 1008–20 (2016).

20. Li, P. et al. Epigenetic dysregulation of enhancers in neurons is associated with Alzheimer’s disease pathology and cognitive symptoms. Nat. Commun. 10, 2246 (2019).

21. Gasparoni, G. et al. DNA methylation analysis on purified neurons and glia dissects age and Alzheimer’s disease-specific changes in the human cortex. Epigenetics Chromatin 11, 41 (2018).

22. Meissner, A. et al. Reduced representation bisulfite sequencing for comparative high-resolution DNA methylation analysis. Nucleic Acids Res. 33, 5868–5877 (2005).

23. Boyle, P. et al. Gel-free multiplexed reduced representation bisulfite sequencing for large-scale DNA methylation profiling. Genome Biol. 13, R92 (2012).

24. The Gene Ontology Resource: 20 years and still GOing strong. Nucleic Acids Res. 47, D330–D338 (2019).

25. Fabregat, A. et al. The Reactome Pathway Knowledgebase. Nucleic Acid Res. 46, 649–655 (2018).

26. Overall, R. W., Paszkowski-Rogacz, M. & Kempermann, G. The Mammalian Adult Neurogenesis Gene Ontology (MANGO) Provides a Structural Framework for Published Information on Genes Regulating Adult Hippocampal Neurogenesis. PLoS One 7, (2012).

27. Koopmans, F., et al. SynGO: An Evidence-Based, Expert-Curated Knowledge Base for the Synapse. Neuron 103, 217–234 (2019).

28. Hadad, N. et al. Caloric restriction mitigates age-associated hippocampal differential CG and non-CG methylation. Neurobiol. Aging 67, 53–66 (2018).

29. Rinaldi, L. et al. Dnmt3a and Dnmt3b Associate with Enhancers to Regulate Human Epidermal Stem Cell Homeostasis. Cell Stem Cell 19, 491–501 (2016).

30. Nan, X., Campoy, F. J. & Bird, A. MeCP2 Is a Transcriptional Repressor with Abundant Binding Sites in Genomic Chromatin. Cell 88, 471–481 (1997).

31. Gabel, H. W. et al. Disruption of DNA-methylation-dependent long gene repression in Rett syndrome. Nature 522, 89–93 (2015).

32. Stroud, H. et al. Early-Life Gene Expression in Neurons Modulates Lasting Epigenetic States. Cell 171, 1151–1154 (2017).

33. Merico, D., Isserlin, R., Stueker, O., Emili, A. & Bader, G. D. Enrichment map: A network-based method for gene-set enrichment visualization and interpretation. PLoS One 5, (2010).

34. Kempermann, G., Kuhn, H. G. & Gage, F. H. Experience-induced neurogenesis in the senescent dentate gyrus. J. Neurosci. 18, 3206–3212 (1998).

35. Su, Y. et al. Neuronal activity modifies the chromatin accessibility landscape in the adult brain. Nat. Neurosci. 20, 476–483 (2017).

36. Mostafavi, S. et al. A molecular network of the aging human brain provides insights into the pathology and cognitive decline of Alzheimer’s disease. Nat. Neurosci. 21, 811–819 (2018).

37. Wingo, A. P. et al. Large-scale proteomic analysis of human brain identifies proteins associated with cognitive trajectory in advanced age. Nat. Commun. 10, 1619 (2019).

38. Maegawa, S. et al. Caloric restriction delays age-related methylation drift. Nat. Commun. 8, (2017).

39. Zhang, T. Y. et al. Environmental enrichment increases transcriptional and epigenetic differentiation between mouse dorsal and ventral dentate gyrus. Nat. Commun. 9, 1–11 (2018).

40. Lewis, J. D. et al. Purification, sequence, and cellular localization of a novel chromosomal protein that binds to Methylated DNA. Cell 69, 905–914 (1992).

41. Gabel, H. W. et al. Disruption of DNA-methylation-dependent long gene repression in Rett syndrome. Nature (2015). doi:10.1038/nature14319

42. Kinde, B., Wu, D. Y., Greenberg, M. E. & Gabel, H. W. DNA methylation in the gene body influences MeCP2-mediated gene repression. Proc. Natl. Acad. Sci. U. S. A. 113, 15114–15119 (2016).

43. Guy, J., Cheval, H., Selfridge, J. & Bird, A. The Role of MeCP2 in the Brain. Annu. Rev. Cell Dev. Biol. 27, 631–652 (2011).

44. Ip, J. P. K., Mellios, N. & Sur, M. Rett syndrome: insights into genetic, molecular and circuit mechanisms. Nat. Rev. Neurosci. 19, 368–382 (2018).

45. Li, H. et al. Cell cycle-linked MeCP2 phosphorylation modulates adult neurogenesis involving the Notch signalling pathway. Nat. Commun. 5, 5601 (2014).

46. Osenberg, S. et al. Activity-dependent aberrations in gene expression and alternative splicing in a mouse model of Rett syndrome. Proc. Natl. Acad. Sci. U. S. A. 115, E5363– E5372 (2018).

47. Stilling, R. M. et al. De-regulation of gene expression and alternative splicing affects distinct cellular pathways in the aging hippocampus. Front. Cell. Neurosci. 1–15 (2014). doi:10.3389/fncel.2014.00373

48. Zhao, B. et al. Somatic LINE-1 retrotransposition in cortical neurons and non-brain tissues of Rett patients and healthy individuals. PLOS Genet. 15, e1008043 (2019).

49. Cardelli, M. The epigenetic alterations of endogenous retroelements in aging. Mech. Ageing Dev. 174, 30–46 (2018).

50. Rampon, C. et al. Effects of environmental enrichment on gene expression in the brain. PNAS 97, 12880–12884 (2000).

51. Zhang, Y. et al. Transcriptomics of Environmental Enrichment Reveals a Role for Retinoic Acid Signaling in Addiction. Front. Mol. Neurosci. 9, (2016).

52. Sahay, A. et al. Increasing adult hippocampal neurogenesis is sufficient to improve pattern separation. Nature 472, 466–470 (2011).

53. Kempermann, G. & Gage, F. H. Experience-Dependent Regulation of Adult Hippocampal Neurogenesis: Effects of Long-Term Stimulation and Stimulus Withdrawal. Hippocampus 9, 321–332 (1999).

54. Ben Abdallah, N. M. B., Slomianka, L., Vyssotski, A. L. & Lipp, H. P. Early age-related changes in adult hippocampal neurogenesis in C57 mice. Neurobiol. Aging 31, 151–161 (2010).

55. Jaeger, B. N. et al. A novel environment-evoked transcriptional signature predicts reactivity in single dentate granuel neurons. Nat. Commun. 9, 3084 (2018).

56. Fernandez-Albert, J. et al. Immediate and deferred epigenomic signatures of in vivo neuronal activation in mouse hippocampus. Nat. Neurosci. 1–13 (2019). doi:10.1038/s41593-019-0476-2

57. Kempermann, G., Gast, D. & Gage, F. H. Neuroplasticity in Old Age: Sustained Fivefold Induction of Hippocampal Neurogenesis by Long-term Environmental Enrichment. Ann. Neurol. 52, 135–143 (2002).

58. Cortese, G. P., Olin, A., O’Riordan, K., Hullinger, R. & Burger, C. Environmental enrichment improves hippocampal function in aged rats by enhancing learning and memory, LTP, and mGluR5-Homer1c activity. Neurobiol. Aging 63, 1–11 (2018).

59. Speisman, R. B. et al. Environmental enrichment restores neurogenesis and rapid acquisition in aged rats. Neurobiol. Aging 34, 263–274 (2013).

60. Krueger, F. & Andrews, S. R. Bismark: a flexible aligner and methylation caller for Bisulfite-Seq applications. Bioinformatics 27, 1571–1572 (2011).

61. Akalin, A. et al. methylKit: a comprehensive R package for the analysis of genome-wide DNA methylation profiles. Genome Biol. 13, R87 (2012).

62. Kinsella, R. J. et al. Ensembl BioMarts: a hub for data retrieval across taxonomic space. Database 2011:bar03, (2011).

63. Pagès, H., Carlson, M., Falcon, S. & Li, N. AnnotationDbi: Annotation Database Interface. R package version 1.42.1. (2018).

64. Karolchik, D. et al. The UCSC Table Browser data retrieval tool. Nucleic Acids Res. 32, 493–496 (2004).

65. Zerbino, D. R., Wilder, S. P., Johnson, N., Juettemann, T. & Flicek, P. R. The Ensembl Regulatory Build. Genome Biol. 16, 1–8 (2015).

66. Khan, A. & Zhang, X. dbSUPER: a database of super-enhancers in mouse and human genome. Nucleic Acids Res. 44, D164–71 (2016).

67. Lawrence, M., et al. Software for Computing and Annotating Genomic Ranges. PLOS Comput. Biol. 9, 1–10 (2013).

68. Shannon, P. & Richards, M. MotifDb: An Annotated Collection of Protein-DNA Binding Sequence Motifs. R Packag. version 1.26.0. (2019).

69. Mclean, C. Y. et al. GREAT improves functional interpretation of cis -regulatory regions. Nat. Biotechnol. 28, 495–501 (2010).

70. Yu, G. & He, Q. Molecular BioSystems ReactomePA: an R / Bioconductor package for reactome pathway analysis and visualization. Mol. Biosyst. 12, 477–479 (2016).

71. Stubbs, T. M. et al. Multi-tissue DNA methylation age predictor in mouse. Genome Biol. 18, (2017).

72. Thompson, M. J. et al. A multi-tissue full lifespan epigenetic clock for mice. Aging (Albany. NY). 10, 2832–2854 (2018).

